# Structure of the NuA4 acetyltransferase complex bound to the nucleosome

**DOI:** 10.1101/2022.07.12.499507

**Authors:** Keke Qu, Kangjing Chen, Hao Wang, Xueming Li, Zhucheng Chen

## Abstract

DNA in eukaryotes wraps around the histone octamer to form nucleosomes^1^, the fundamental unit of chromatin. The N-termini of histone H4 interact with nearby nucleosomes, and play an important role in the formation of high order chromatin structure and heterochromatin silencing^2-4^. NuA4 in yeast and its homolog Tip60 complex in mammalian cells are the key enzymes that catalyze H4 acetylation, which in turn regulate chromatin packaging, and function in transcription activation and DNA repair^5-10^. Here we report the cryo-EM structure of NuA4 from *Saccharomyces cerevisiae* bound to the nucleosome. NuA4 comprises two major modules: the catalytic histone acetyltransferase (HAT) module and the transcription activator-binding TRA module. The nucleosome is mainly bound by the HAT module, and positioned close to a polybasic surface of the TRA module, which is important for the optimal activity of NuA4. The nucleosomal linker DNA carrying the upstream activation sequence is oriented towards the conserved, transcription-activator-binding surface of the Tra1 subunit, which suggests a potential mechanism of NuA4 to act as a transcription co-activator. The HAT module recognizes the disk face of the nucleosome through the H2A-H2B acidic patch and the nucleosomal DNA, projecting the catalytic pocket of Esa1 to the N-terminal tail of H4, supporting its function in selective acetylation of H4. Together, our findings illustrate how NuA4 is assembled, and provide mechanistic insights into nucleosome recognition and transcription coactivation by a histone acetyltransferase.

---

NuA4 contains 13 subunits, of which Esa1 is the catalytic subunit for the acetylation reaction, and Tra1 is the scaffold subunit for binding to a variety of transcription activators/factors (TFs) to recruit NuA4 for targeted gene activation^11-13^. Esa1 interacts with Epl1, Yng2 and Eaf6 to form a Piccolo subcomplex^14^, which achieves selective H4 acetylation in the context of the nucleosome through a position-based mechanism^15^. Interestingly, Tra1 is the common subunit shared by another transcription co-activator HAT complex, SAGA, which catalyzes H3 acetylation on nucleosomes^12,13^. Despite the fundamental importance, currently available structures of the NuA4 complex are at very low resolutions^16-18^, and it remains unclear how the HAT complexes interact with the nucleosome substrate^15,19^.

## Structural determination of the complex

To elucidate the mechanism of NuA4, we determined the cryo-EM structure of NuA4 bound to the nucleosome. Endogenous NuA4 complex was purified from yeast cells (Extended Data Fig. 1a), mixed with nucleosome core particles (NCPs), and subjected to single particle cryo-EM analyses (Extended Data Fig. 2a). A low-resolution structure was achieved, which nevertheless revealed the molecular envelop of the large subunit Tra1 and a nucleosome-like density nearby (Extended Data Fig. 2b). To reduce the flexibility of the complex, the upstream activation sequence (UAS) for Gal4 binding was engineered into the linker DNA sequence of the nucleosome (UAS-20N20), and the transcription activator Gal4-VP16 was added to the NuA4-NCP complex. It is interesting to notice that the UAS placed at different positions promoted the HAT activity of NuA4 (Extended Data Fig. 1b), suggesting plasticity of the complex. In the presence of Gal4-VP16, the structure was determined at an overall resolution of ∼8.7 Å, and revealed the envelop of the nucleosome, and an unknown density on the disk face (Extended Data Fig. 2c-2d). Focused refinements did not resolve the unknown density.

Considering that the bisubstrate inhibitor, carboxymethyl coenzyme A (CMC), stably binds to the Esa1 catalytic pocket^20,21^, and the chromo domain of Eaf3 recognizes methylated H3K36^22,23^, we modified the histones at H4K16 to contain CMC, and H3 to contain trimethylated Lys36 to further restrain the flexibility of the complex (Extended Data Fig. 3). Under these conditions, the structure was determined at an overall resolution similar as above, and focused refinements led to resolutions of 3.1 and 3.4 Å at the TRA module and the HAT module bound on the top of the nucleosome, respectively, with local resolutions of 2.7-2.8 Å (Fig. 1, Extended Data Figs. 4 and 5, Extended Data Table 1). Whereas the linker DNA in the high-resolution map is barely detected because of the focused classification and refinement, the overall map of the holo-complex covered ∼25 bp linker DNA, and a local map (at a resolution of ∼6.7 Å) covered a longer linker DNA (∼30 bp) and helped to orientate the UAS-20N20 nucleosome (Fig. 1a and Extended Data Figs. 4b). The orientation of the nucleosome was also supported by the high-resolution maps of the central base pairs (Extended Data Fig. 5b). The subunit interactions were confirmed by cross-linking mass-spectrometry (CL-MS, Extended Data Fig. 1c and 1d).

**Figure 1.**
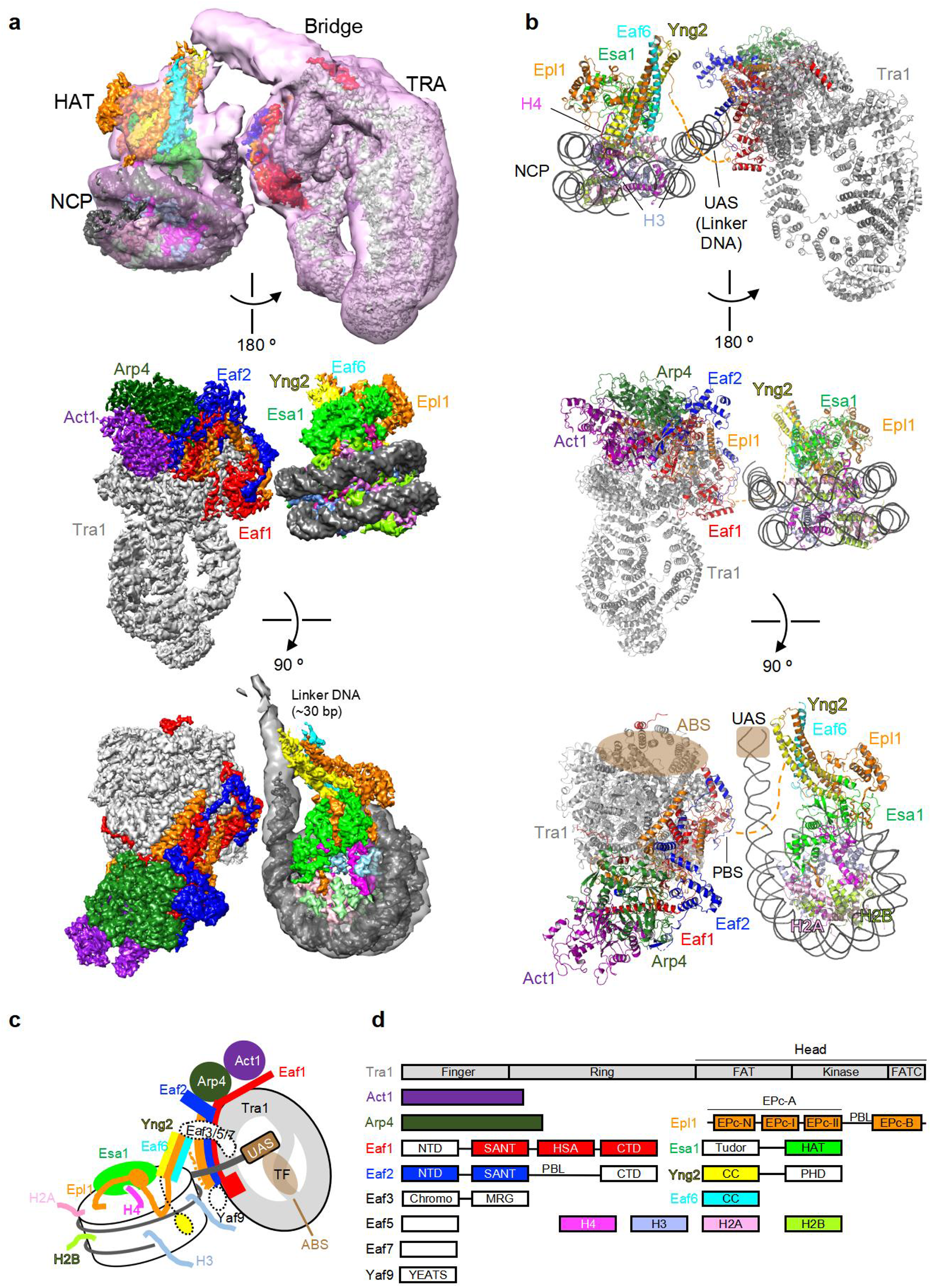
Overall structure of the NuA4 complex bound to the nucleosome. (**a**) Three different views of the high-resolution composite maps of the NuA4-NCP complex. The low-resolution molecular envelops of the all-overall complex (top panel) and the UAS linker DNA (bottom panel) are shown. (**b**) Three views of the ribbon model of the NuA4-NCP complex. The linker DNA of 25 bp is shown, and the last 5 bp DNA is colored as “UAS”. (**c**) Model of the assembly and mechanism of nucleosome recognition of NuA4. TF is schematically illustrated to interact with the ABS and recognize the UAS. (**d**) Domain organization of the NuA4 subunits and histones. Proteins are color coded, with those not structurally resolved in white. CC, coiled coil; PBL, polybasic loop.

## Overall structure of NuA4

NuA4 is organized into two major modules: the catalytic HAT module and the transcription activator-binding TRA module (Fig. 1a). The HAT module of NuA4 is essentially the core Piccolo subcomplex^14^, composed of the catalytic subunit Esa1, the N-terminal EPc-A fragment of Epl1, the PHD domain-containing Yng2, and Eaf6 (Fig. 1b-d). The TRA module includes the largest subunit Tra1, Actin, Arp4, Eaf2, Eaf1, and the EPc-B domain of Epl1. Actin, Arp4, and the HSA domain of Eaf1 assemble into a submodule (referred as to the ARP submodule), and bind to the head of Tra1 as described previously^17^. The two major modules are linked through a disordered connecting loop between the EPc-A and EPc-B domains of Epl1, consistent with the association of Piccolo to the rest of NuA4 through the Epl1 C-terminal domain^14^. Eaf3, Eaf5, and Eaf7 form a subcomplex^24^, but their structure could not be clearly resolved. Eaf3 extensively crosslinked to the connecting loop of Epl1 (Extended Data Fig. 1c), suggesting that the bridge density connecting the HAT and TRA modules resulted from the Eaf3/5/7 subcomplex (Fig. 1a). Multi-body refinement analysis indicated different modes of relative movement of the two major modules of NuA4 around a polybasic surface (PBS) formed by the Epl1 and Eaf2 subunits of the TRA module, explaining the plasticity of the complex and the difficulty in resolving a high-resolution structure.

## Activator binding surface of Tra1

The structure of the complex provides the framework for NuA4 to function as a transcription coactivator. Tra1 comprises > 3,000 residues, binds to various TFs and recruits NuA4 for targeted gene expression^13^. The linker DNA carrying the UAS packs against the edge of the TRA module, and projects toward the Finger and FAT domains of Tra1 (Fig. 2a). Intriguingly, the exposed surface of Tra1 facing the UAS is highly conserved from yeast to humans, suggesting functional importance, whereas the surface at the opposite side is more variable (Fig. 2b). We propose that the conserved surface of Tra1 serves as the activator binding surface (ABS). In support of this notion, His400, which is required for Gal4 binding, maps to the conserved surface of the Finger domain^25^. Furthermore, the Gcn4 and Rap1 binding sites^26^, residues 88-165 and 319-399, also map to the nearby region. The VP16 binding site is less well-defined, and was proposed to involve the Tra1 C-terminal domain^27^. The structure of VP16 was not resolved currently. Consistent with the proximity of linker DNA to the FAT domain of Tra1 (Fig. 2a), Gal4-VP16 was crosslinked to multiple residues (Lys2808, Lys2984, and Lys3161) of the FAT domain of Tra1 (Extended Data Fig. 1c). The conserved surface extends along the Finger domain to the FAT domain, spans a large distance of over 100 Å, and potentially interacts with different TFs, which independently and/or cooperatively recognize the adjacent linker DNA (UAS) of the nucleosome to achieve targeted recruitment of NuA4 for gene activation.

**Figure 2.**
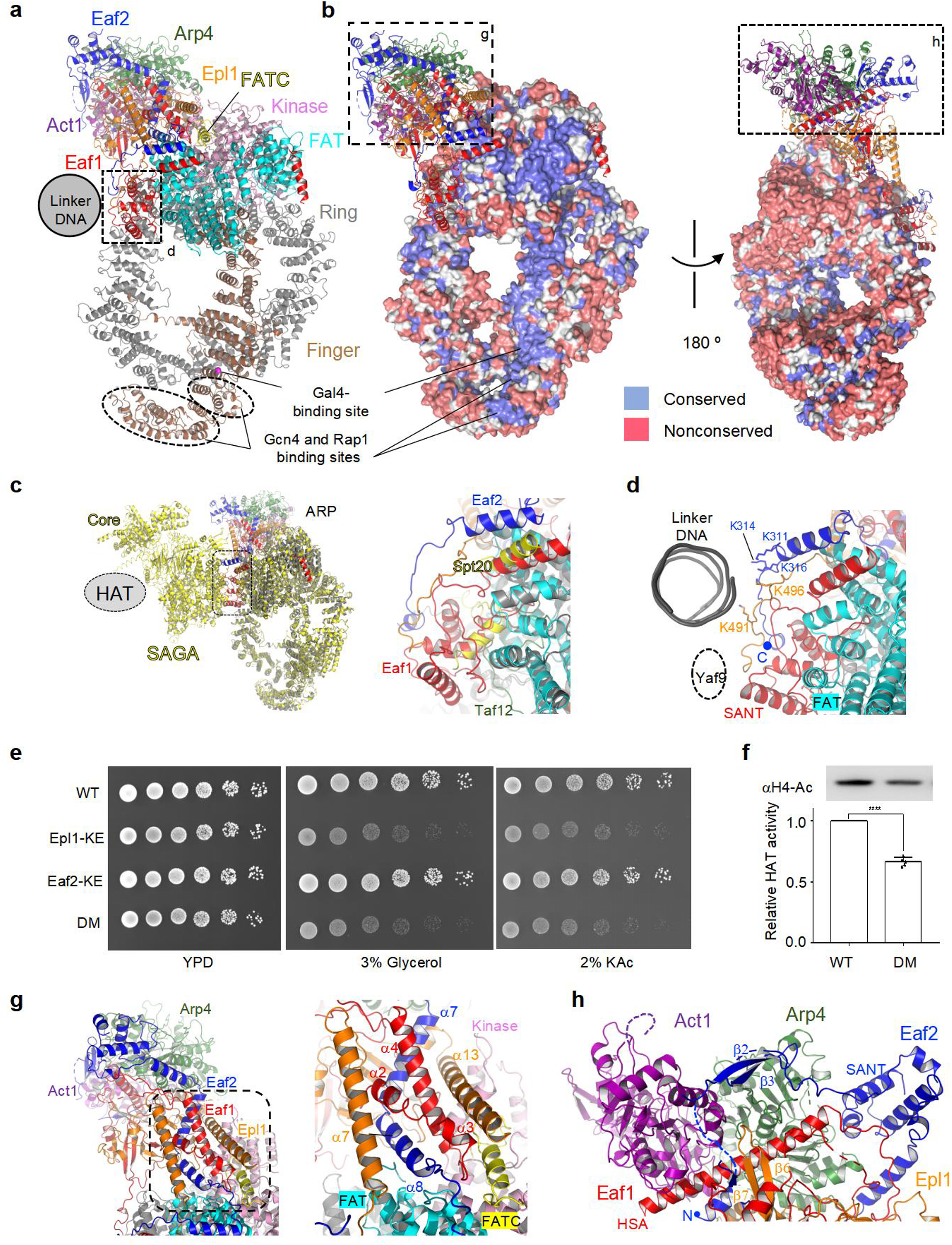
Structure of the TRA module. (**a**) The ribbon model of the TRA module with the five Tra1 domains Ring, Finger, FAT, Kinase, and FATC colored differently. The other subunits are colored as in Fig. 1. The boxed region is enlarged for analysis in (d). (**b**) Surface conservation of Tra1. Increasing conservation scores of the residues are shown in the red-white-blue spectrum^39^. The boxed regions are enlarged for analysis in (g) and (h). (**c**) Structural comparison of NuA4 and SAGA (colored yellow, PDB code 6T9I)^19^. The structures of Tra1 are aligned. The position of the HAT module of SAGA is schematically illustrated, and the boxed region is enlarged for analysis on the right. (**d**) Structure of the PBS region of the TRA module. (**e**) Growth phenotype of the WT and three mutant yeast cells under indicated conditions. KAc, potassium acetate. Three independent sets of experiments were performed, and the presentative one was showed. (**f**) Relative HAT activity of the purified WT and DM mutant NuA4 complex. The activity of the WT sample was normalized to 1. Error bars indicate SD (n = 5 technical replicates). ****, p<0.0001. (**g**) Structure of the ARP submodule. The boxed region is enlarged for analysis on the right. (**h)** Interactions between the Act1-Arp4 dimer with Eaf1, Eaf2 and Epl1.

Tra1 is also responsible for transcription activator binding in the SAGA complex^12,13^. Interestingly, the Eaf1-binding surface of Tra1 in NuA4 overlaps with the Spt20-Taf12 binding surface in the SAGA complex^19^ (Fig. 2c), explaining the mutual exclusive assembly of Tra1 into NuA4 and SAGA^28^. The HAT module in SAGA is oriented relative to Tra1 in a manner analogous to that of NuA4, suggesting that the ABS of Tra1 identified in NuA4 also provides the mechanism of nucleosome recruitment for SAGA.

## Polybasic surface of Epl1 and Eaf2

The PBS of the TRA module is composed of polybasic loops of Eaf2 and Epl1 (Fig. 2d), and positioned adjacent to the linker DNA at the nucleosome edge (Fig. 1b). Lys311, Lys314 and Lys316 of Eaf2 are highly conserved (Extended Data Fig. 6), and exposed to the nucleosomal linker DNA. Likewise, Lys491 and Lys496 of Epl1 are exposed, and polybasic sequences at the equivalent positions are found in the homologs (Extended Data Fig. 7). There is a gap between the PBS and the linker DNA, suggesting long-range charge-charge interactions. Given the flexibility between the TRA and HAT modules, it is very likely that the PBS interacts with varied positions of the DNA, in line with HAT activation by the UAS located at different positions of the linker DNA (Extended data Fig. 1b).

To examine the importance of the PBS, we replaced the wild type (WT) Epl1 with the polybasic loop mutant (K491E/K496E, Epl1-KE for simplicity). Similarly, we introduced the mutant Eaf2 (K311E/K314E/K316E, Eaf2-KE), and combination of the Epl1 and Eaf2 mutations in the double mutant (DM). Whereas the mutant cells showed normal growth comparable to the WT cells in YPD, the Epl1-KE mutant showed growth defect using alternative carbon sources, such as glycerol and acetate, other than dextrose (Fig. 2e). The Eaf2-KE mutant did not show obvious defect, and the DM mutant displayed a slow growth phenotype similar to the Epl1-KE mutant. The data indicate that the in vivo function of NuA4 depends on the PBS, the polybasic loop of Epl1 in particular, which is in closer to the linker DNA (Fig. 2d).

Consistent with the data in cells and the structure, the DM mutations did not perturb the integrity of the complex (Extended Data Fig. 1e), but the purified enzyme showed a reduced HAT activity relative to the WT NuA4 complex in vitro (Fig. 2f and Extended Data Fig. 1e). Together, the data indicate that the interaction of the nucleosomal linker DNA with the PBS is important for the optimal activity of NuA4.

The polybasic loops of Eaf2 and Epl1 bind to the SANT domain of Eaf1 (Fig. 2d), which was previously assigned to a position close to the ARP submodule based on the low-resolution map^17^. However, our high-resolution map indicates that the Eaf1 SANT domain interacts with the FAT domain of Tra1 (Extended Data Fig. 5a), consistent with the crosslinking analysis (Extended Data Fig. 1c) and the previous studies showing that this domain is required for Tra1 association^29^. The C-terminal domain (CTD) of Eaf2 following the polybasic loop is disordered. Given that the YEATS domain-containing subunit Yaf9 crosslinked to the Eaf2 CTD^30^ (Extended Data Fig. 1c), the structure suggests that Yaf9 is tethered close to the nucleosome around the PBS of the TRA module, consistent with its role in binding to the acetylated H3 tails through the YEATS domain^31^.

## Structure of the ARP submodule

The ARP submodule binds to the head of Tra1 through a helical base formed by Eaf1, Eaf2 and Epl1 (Fig. 2b and 2g). The structure explains the multiple roles of Eaf1 in the NuA4 assembly^18,29,32^. The SANT domain of Eaf1 binds to the edge of Tra1 (Fig. 2d), and the HSA helix interacts with the Act1-Arp4 dimer^17^ (Fig. 2h). The α2-α4 helices of Eaf1 (residues 285-340, Extended Data Fig. 8) bundle with α7-α8 of Eaf2 and α13 of Epl1, forming the helical base that binds to the surfaces of the FAT, kinase, and FATC domains of Tra1 (Fig. 2g). This structure is consistent with the importance of a fragment of Eaf1 before the HSA helix for Tra1 binding^18^. The structure is also consistent with the previous studies showing that deletion of the FATC domain (residues 3711-3724) of Tra1 causes post activator-recruitment defects in transcription^26^, which likely perturbs the binding of the ARP submodule.

Actin and Arp4 in NuA4, previously proposed to function in substrate recruitment^17,33^, locate at a position proximal to, but not directly contacting the nucleosome (Fig. 1b), suggesting that their function in substrate recruitment is probably achieved in an indirect manner. The Act1-Arp4 heterodimer in NuA4 is stabilized by a series of previously unrecognized interactions (Fig. 2h). A highly conserved β-hairpin (β6-β7) of Epl1 interacts with the HSA helix and the Act1-Arp4 heterodimer (Extended Data Figs. 5a and 7). Likewise, an N-terminal β-hairpin (β2-β3) of Eaf2 interacts with Actin and Arp4, which may play a role analogous to that of the β-hairpin of Rtt102 in stabilizing Arp7-Arp9 in the SWI/SNF complex^34,35^. Moreover, Arp4 packs against the SANT domain of Eaf2, the density of which was assigned to the Eaf1 SANT domain^17^. The high-resolution map clearly supports the sequence identity of Eaf2 (Extended Data Fig. 5a). Our findings reveal the extensive contacts between Eaf2 and the Act1-Arp4 dimer, which provide the structural basis for them to form a stable submodule shared by the NuA4 and SWR1 complexes^11^.

## Nucleosome recognition of the HAT module

The HAT module of NuA4 is organized into two parts, the catalytic core formed by Esa1 and the EPc-I domain of Epl1, and the helical bundle formed by the EPc-II domain of Epl1, Eaf6 and Yng2 (Fig. 3a). This structure largely maintains the structure of the isolated Piccolo subcomplex^15^, with the β-hairpin of Esa1 and the helical bundle tilting upward to avoid clashing with the bound nucleosome (Extended Data Fig. 9a). Importantly, similar to Piccolo^15^, the HAT module of the holoenzyme binds to the nucleosome disk face, with the catalytic pocket of Esa1 oriented towards the H4 N-terminus, which shows notable EM density extending from the nucleosome into the catalytic center (Extended Data Fig. 5b). The CMC-modified H4K16 binds to the acetyl-CoA-binding pocket of Esa1 at a position similar to those observed before^15,21^ (Extended Data Fig. 9b and 9c). The positioning of the catalytic pocket above the H4 tail is not an artifact caused by the H4-modification, because a similar spatial arrangement was observed for the unmodified nucleosome (Extended Data Fig. 2d). Therefore, the modular nature of NuA4 ensures that the holoenzyme retains the position-based mechanism of the catalytic Piccolo subcomplex for H4 recognition^15^.

**Figure 3.**
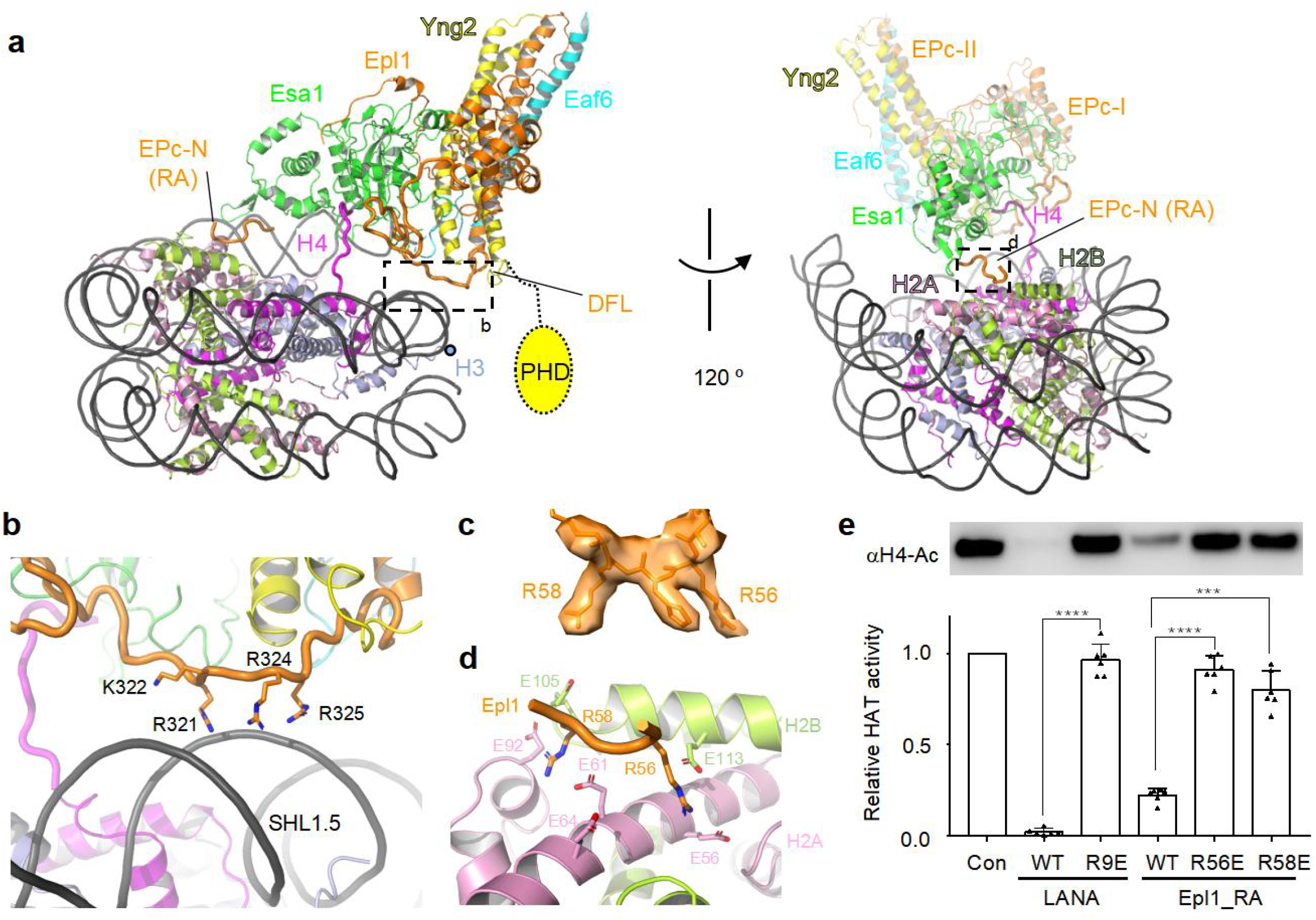
Structure of the HAT module bound to the nucleosome. (**a**) Two views of the structure of the HAT module bound to the disk face of the nucleosome. RA, arginine anchor. The boxed regions are enlarged for analysis in (b) and (d). (**b**) Binding of the DFL to the nucleosome DNA. (**c**) Local EM density of the arginine anchors. (**d**) Interaction of the arginine anchors of NuA4 with the H2A-H2B acidic patch. (**e**) Relative HAT activity of NuA4 to the H4 of the nucleosome in the absence and presence of the LANA and arginine anchor peptides. Error bars indicate SD (n = 6 technical replicates). ***, p=0.0001; ****, p<0.0001.

The HAT module binds to the nucleosome disk face through two elements of Epl1, the double function loop (DFL)^15^ and the arginine anchors (Fig. 3a). The DFL connects the helical bundle with the catalytic core, contains multiple positively charged residues, including Arg321, Lys322, Arg324, and Arg325, and is in close proximity to the minor grove of the DNA at the super-helical location 1.5 (SHL1.5) of the nucleosome (Fig. 3b and Extended Data Fig. 5b). In support of the functional importance of the DFL, these residues are highly conserved across evolution (Extended Data Fig. 7), and the mutations of Lys322 and Arg322 diminish the HAT activity of Piccolo towards the nucleosome in vitro, and result in sensitivity to DNA damage in vivo^15^.

A basic region of the EPc-N domain of Epl1 binds to the nucleosome^36^. The high-resolution map resolves two arginine anchor residues, Arg56 and Arg58, of the EPc-N domain, which interact with the acidic patch of H2A-H2B (Fig. 3c and 3d). Consistent with the structure, Lys59 of Epl1 was crosslinked to Lys95 of H2A (Extended Data Fig. 1c). The importance of acidic patch recognition was demonstrated by adding WT LANA peptide to compete with NuA4 for binding to the acidic patch^37^.WT LANA peptide markedly reduced the HAT activity of NuA4, whereas the binding-defective mutant peptide did not perturb the activity (Fig. 3e). Likewise, addition of the Epl1 arginine anchor peptide to the HAT assays inhibited NuA4, whereas addition of the R56E or R58E mutant peptides largely abolished the inhibition. Together, the data indicate that the DFL and arginine anchors of NuA4 recognize two epitopes on the disk face of the nucleosome, and this recognition directs the catalytic pocket of Esa1 towards the H4 N-terminal tail, providing the structural basis for the H4 preference of NuA4.

In summary, our findings provide an integrated model of nucleosome recognition by NuA4 through the coordination of multiple elements, including the ABS and PBS of the TRA module, and the DFL and arginine anchors of the HAT module (Fig. 1c). The PBS and ABS interact with the linker DNA, directly and indirectly, and recruit the nucleosome to the edge of the TRA module, such that the closely linked HAT module binds to the disk face of the nucleosome to achieve selective acetylation of the H4 tails. Moreover, the tethering histone-tail-binding domains, such as the PHD domain of Yng2 and the YEATS domain of Yaf9, are spatially arranged around the nucleosome, allowing NuA4 to read histone modification signals^31,38^. The relative orientation of the two major modules of NuA4 shows a large degree of plasticity, which may nevertheless provide the adaptability required in response to different transcription activators and chromatin cues. All the subunits except Eaf5 of yeast NuA4 are conserved in the Tip60 complex in human cells, suggesting that our structure provides a good model for human Tip60.

## METHODS

### Preparation and purification of native NuA4 complex

The yeast strain BJ2168 (a gift from Soo-Chen Cheng lab) used in this study carries a tandem His-TEV-protein A tag at the C-terminus of Eaf1 subunit. The homology recombination cassettes were amplified by PCR from the modified plasmid pFA6a-TAP-KanMX6. The PCR products were transformed into *Saccharomyces cerevisiae* cells by lithium acetate method and selected on G418-YPD solid medium^40^. Integration of the cassettes was confirmed by PCR and Western blot.

To prepare NuA4, yeast cells were cultured in YPD medium at 30 °C to an OD_600_ of 4. Cells were pelleted, washed, and resuspended in the lysis buffer (50 mM HEPES, 500 mM KCl, 2 mM MgCl_2_, 20% glycerol and protease inhibitors cocktail, pH 7.6). The cell suspension was dripped into liquid nitrogen and then ground into powder using a chilled pestle. The powder was thawed at room temperature and centrifuged first at 16,000 g for 1 hour and the supernatant was centrifuged again at 150,000 g for 1 hour. The supernatant was loaded onto a HisTrap column (GE healthcare), washed by the lysis buffer with 20 mM imidazole and eluted with 300 mM imidazole. The elution was incubated with 1ml IgG Sepharose 6 Fast Flow resin (GE healthcare) overnight, and then cleaved by TEV protease at 18 °C for 1 hour in a buffer containing 20 mM HEPES, 150 mM KCl, 2 mM MgCl_2_, 0.1% NP40, 10% glycerol, pH 7.6. The eluted protein was concentrated and applied to a glycerol gradient centrifugation. Fractions containing NuA4 were pooled, and concentrated to ∼300 nM.

To prepare the sample for EM analysis, NuA4 was mixed with 600 nM Gal4-VP16. The mixture was dialyzed to the binding buffer containing 20 mM HEPES, 50 mM KCl, 2 mM MgCl_2_, 10% glycerol, pH 7.6 for 6 hours. Then 40N20 NCP (900 nM) was added, and incubated on ice for 1 hour. The mixture was cross-linked and purified using Grafix method^41^. The glycerol gradient was prepared using the light buffer containing 10% (v/v) glycerol, 20 mM HEPES, 50 mM KCl, 2 mM MgCl_2_, pH 7.6, and the heavy buffer containing 40% (v/v) glycerol, 20 mM HEPES, 50 mM KCl, 2 mM MgCl_2_, 0.2% glutaraldehyde, pH 7.6. The sample was centrifuged at 38,000 rpm for 16 h at 4 °C using a Beckman SW41 Ti rotor. Fractions containing NuA4-Gal4-VP16-NCP were confirmed by 4.5% native PAGE, stained by SYBR gold. Peak fractions were pooled and dialyzed in the EM buffer containing 20 mM HEPES, 50 mM KCl, 2 mM MgCl_2_, 0.005% NP40, pH7.6 for 4 hours to remove glycerol and concentrated for EM analysis.

### Purification of Gal4-VP16 fusion protein

The Gal4 DNA binding domain (residues 1-147) was cloned from yeast genomic DNA. VP16 activator domain (residues 413-490) was amplified by primer walking. The two fragments were integrated to a pET28b vector by seamless cloning method. *E. coli* Rosetta (DE3) was used for protein expression. Briefly, cells were grown at 37 °C to an OD_600_ of 0.6, and induced by addition of 0.5 mM IPTG. Cells were further cultured for 3 hours at 37 °C and harvested. Cells were lysed by sonication in the lysis buffer (50 mM Tris, 500 mM NaCl, 10% glycerol, 1 mM PMSF, 2 mM Benzamidine, pH 8.0), and the soluble extract was loaded onto Ni^2+^-NTA-agarose resin. The eluted sample was diluted to 50 mM NaCl, loaded onto a resource 15S column and resolved with a linear gradient of 0.05-1 M NaCl. The peak fractions were pooled, concentrated and applied to a Superdex-200 column.

### Preparation of histone H3 modified with methyl-lysine analog

To generate the K36-trimethylated histone H3 variant, double mutants K36C and C110A were introduced into the H3 sequence by site-directed mutagenesis. The tri-methyl-aminoethyl group was introduced to the thiol group of K36C similarly as described^42^. The final product was confirmed by Q-TOF mass spectrometry.

### Preparation of histone H4 modified with CMC

The chemically synthesized H4 peptide (residues 11-22) modified with CMC at K16 was purchased from Scilight-peptide company. H4 (residues R23C-102) was amplified with PCR using WT template and integrated to a pET3c vector by seamless cloning method. Expression and purification of the truncated histone was performed similarly as the WT, with some modifications. Protein was expressed in *E. coli* Rosetta (DE3) cells. The histone was dialyzed once against 5 mM DTT solution, and at least twice against 0.05% TFA solution after the ion exchange purification step. Removal of the N-terminal methionine was confirmed by MALDI-TOF mass spectrometry.

The final product, histone H4 modified with CMC, was assembled by native chemical ligation as similarly described^43,44^, and confirmed by Q-TOF mass spectrometry. First, H4 (residues 11-22) K16CMC (1.0 equiv.) was dissolved in the aqueous phosphate (0.2 M) buffer containing 6 M guanidinium chloride. 1 M NaNO_2_ aqueous solution (5 equiv.) was added drop-wise and stirred for 20 min at −12 °C to produce a peptide acyl azide. MPAA (4-mercaptophenylacetic acid, 50 equiv.) was added and the pH value was adjusted to 6.3. The reaction was then removed to the room temperature and kept for 5 min. H4 (residues R23C-102) (1.2 equiv.) was added and the reaction buffer was adjusted to pH 6.5 with 2 M NaOH to initiate native chemical ligation. The reaction mixture was stirred at 18 °C overnight and reduced by TCEP (50 mM, pH 7.0). The final reaction mixture was purified by a resource 15S column. The peak fractions were pooled, lyophilized and stored at −80 °C.

### Preparation of nucleosome

Wild type Xenopus laevis histones H2A, H2B, H3 and H4 were expressed and purified as described^45^. For octamer preparation, histones H2A, H2B, H3 (or H3K36me3) and H4 (or H4K16-CMC) were mixed at a molar ratio of 1:1:1:1 and dialyzed three times against the refolding buffer (20 mM Tris, 2 M NaCl, 10 mM DTT, pH7.5). The sample was concentrated and applied to a Superdex 200 size exclusion column. Peak fractions were pooled, concentrated and frozen in liquid nitrogen.

The 40N20 (UAS-20N20) DNA fragment (**TCCGGAGGACTGTCCTCCGG**GGACCCTATACGCGGCCGCC<u>ATCGAGAAT CCCGGTGCCGAGGCCGCTCAATTGGTCGTAGACAGCTCTAGCACCGCTTAA ACGCACGTACGCGCTGTCCCCCGCGTTTTAACCGCCAAGGGGATTACTCCCTAGTCTCCAGGCACGTGTCAGATATATACATCCGATAGCTTGTCGAGAAGTA CTAG, the Gal4 DNA binding sequence in bold, and the “601” sequence underlined) was prepared by PCR, using the Widom 601 sequence as the template. The 5′ variant linker DNA sequences used as follows:</u>

UAS-0N20 (**TCCGGAGGACTGTCCTCCGG**)

UAS-10N20 (**TCCGGAGGACTGTCCTCCGG**TCGTGCCTGT)

con-20N20 (GAGTTCATCCCTTATGTGATGGACCCTATACGCGGCCGCC)

The PCR product was purified by anion-exchange chromatography and concentrated. Nucleosome reconstitution was performed as described^45^.

### Electron Microscopy

The NuA4 complex was first studied by negative stain EM. Briefly, 4 µl of sample was loaded onto a glow-discharged copper grid coated with a thin carbon film. The sample was incubated for 1 min and grids were blotted and immersed in uranyl acetate (2%, w/v) for negative stain.

For cryo-EM analysis, 4 µl of sample was applied to Quantifoil R2/1 Au 200 mesh grids which were glow-discharged for 25 sec, and subsequently incubated for 60 sec before blotting and vitrification by plunging into liquid ethane with a Vitrobot (FEI) operated at 8 °C and 100% humidity.

### Negative stain data collection and image processing

In the negative stain data collection, the NuA4-NCP samples were observed using a 200 kV Tecnai F20 microscope (FEI) equipped with a Gatan Ultrascan 4000 camera at a magnification of 62,000×, corresponding to pixel size of 1.35 Å on the images. Defocus ranging from −1.0 to −2.5 µm and a total dose of ∼40 e^-^/Å^2^ were used. The defocus parameters were estimated by CTFFIND3^46^. A total of 10,354 particles were picked using the ‘e2boxer.py’ subroutine in the EMAN2 suit^47^. Two-dimensional (2D) classification was carried out in Relion 1.4 to remove the bad particles^48^. A sphere generated by SPIDER was used as the initial model for the first round of three-dimensional (3D) classification^49^. After several rounds of three-dimensional classification, 8,739 particles were selected, and subjected to the final three-dimensional reconstruction.

### Cryo-EM data collection and image processing

The dataset of NuA4-Gal4-VP16-NCP (H4 modified) was acquired on a Thermo Fisher Scientific Krios G3i operated at 300 kV, equipped with a K3 direct electron detector and GIF Quantum energy filter (Gatan). The movie stacks were automatically recorded using AutoEMation2 in super-resolution mode^50^, at a magnification of 81,000× for a final pixel size of 0.54125 Å/pixel in the super-resolution mode with the defocus values ranging from −1.3 to −1.8 µm. The total electron dose was 50 e^-^/ Å^2^ fractionated in 32 frames (exposure time 2.56 sec).

A total of 11,000 dose-fractionated image stacks were aligned using MotionCor2 with twofold binning^51^, resulting in a pixel size of 1.0825 Å/pixel. CTF parameters were estimated using CTFFIND4^52^. Particle picking, two-dimensional (2D) classification and three-dimensional (3D) classification were carried out in Relion3.0^48^. Particles were extracted using a box size of 376 × 376 pixels, and normalized. Reference-free 2D classification and global 3D classification with a NuA4-NCP map derived from the NuA4-Gal4-VP16-40N20 dataset low passed to 40 Å as the initial model were performed to screen for “good” particles in the dataset. All “good” particles with fourfold binning were used for 3D refinement with local search resulting in a map at 8.8 Å. Furthermore, these particles were re-extracted without binning and used for global 3D refinement resulting in a map at 4.0 Å. To improve the map of the TRA module, focused 3D classification without image alignment was performed using a soft mask. The class that showed the best density for the TRA module was subjected to another round of focused 3D refinement resulting in a resolution of 3.0 Å. Focused refinement further improved the resolution to 2.7 Å for the ARP submodule and around the Head region of Tra1. To improve the bottom part of Tra1, a soft mask was applied to this region and the following 3D classification without image alignment led to an intact TRA module. 3D refinement with a soft mask on the TRA module led to a map at 3.1 Å.

For reconstruction of the NCP-HAT region, the map of the NCP-HAT module low passed to 40 Å was used as the reference to obtain an improved initial reference. All classes containing the intact NCP-HAT density were re-extracted without binning and used for a global refinement resulting in a map at 7.7 Å resolution. Focused 3D classification without image alignment, and focused refinement led to a resolution at 3.4 Å. For the NCP-arginine-anchor region, all classes showed clear features were combined. Focused 3D classification without image alignment and following 3D refinement led to a resolution at 2.8 Å. For the reconstruction of the HAT-NCP module with long linker DNA, all classes containing the intact HAT-NCP density were re-extracted with twofold binning, used for multi-round of 3D classifications, and for 3D refinement with signal subtraction, which led to a resolution at 6.7 Å.

The different modes of relative movement of the TRA module and the HAT module were carried out by multi-body refinement^53^. It reveals a rotational motion of the TRA module bound to the nucleosome. The first 3 eigenvectors shown in the movie explain 40.4 % of the variance in the data, with 15.2% of variance for component 1, 12.7% of variance for component 2, and 12.5% of variance for component 3. They represent the principal motions in the particles.

The datasets of NuA4-NCP and NuA4-Gal4-VP16-NCP were acquired on an FEI Titan Krios operating at 300 kV with a nominal magnification of 22,500×. Images were recorded by a Gatan K2 Summit detector (Gatan company) with the super-resolution mode, and binned to a pixel size of 1.306 Å. UCSFImage4 was used for data collection under a defocus values ranged from −1.4 to −1.8 µm^54^. Each micrograph was dose-fractionated to 32 frames with 0.25 s exposure time in each frame and the total electron dose was 50 e^-^/Å^2^. Motion correction, CTF estimation, particle picking and reference free 2D classification were the same as above.

For the NuA4-NCP sample, “good” particles were processed with global 3D classification using RELION 3.0 and a negative stain reconstruction of the NuA4-NCP as the first reference to obtain an improved initial reference. After several rounds of 3D classification and refinement, only a low resolution map of the intact complex was obtained (Extended Data Fig. 2b).

For NuA4-Gal4-VP16-NCP sample, all “good” particles collected from 2D classification were used for global 3D classification. After several rounds of 3D classification, the class containing intact NuA4-NCP density was used for 3D refinement, and led to a map with resolution at 8.8 Å. All classes containing the intact TRA module were combined and subjected to 3D refinement, leading to a map with resolution at 3.6 Å (Extended Data Fig. 2d), but the nucleosome-bound HAT did not result in high-resolution map.

### Model building

The model of NuA4 was built by fitting the available structures of Tra1, the catalytic Piccolo subcomplex, the Act1-Arp4-HSA^Eaf1^ subcomplex and Esa1-bound CMC ^15,17,21,55^. The rest of the molecules, including Eaf2, Eaf1 and the EPc-B domain of Epl1, were manually built based on the high-resolution maps in Coot. The structures were refined using Phenix with secondary structure constrains^56^. The asymmetric cryoEM density of the linker DNA, which contains DNA of >25 bp at one side, unambiguously orientates the 40N20 nucleosome into the map. The orientation of the nucleosome is also supported by the high-resolution map of the nucleosome (at a resolution of 2.8 Å), the four central base pairs of which are showed in the Extended Data Fig. 5b.

### Cross-linking and mass spectrometry

Sample preparation was the same as that for the cryo-EM analysis. The mixture was dialyzed to the binding buffer for 4 hours, and crosslinked by 3 mM BS3 for 30 min at room temperature. The reaction was quenched with 50 mM Tris, pH 7.5. The cross-linked sample was applied to a 10-40% glycerol gradient as described above. The protein containing fractions were pooled for mass spectrometry^57^. The data were analyzed with xiVIEW^58^ and displayed with chimeraX^59^.

### Yeast genetics

To generate Eaf2-KE (K311E/K314E/K316E) mutant strain, the C-terminal of Eaf2 (residues 311-476) was first replaced by URA3, and the yeast cells were selected using SC/-Ura medium. Then URA3 was replaced by the C-terminal of Eaf2 with desired mutations, and the cells were selected using 5-FOA medium (YPD + 5FOA). The Epl1-KE (K491E/K496E) mutant strain was generated the same as described above except the C-terminal of Epl1 (residues 491-832) was replaced by URA3. The DM (double mutant) strain harbors both Eaf2 and Epl1 mutations was generated by introducing the C-terminal of Eaf2 mutations into the Epl1-KE mutant strain as described above. The yeast strain BY4741 (MATα *leu2 ura3 his3 met15*) was used, and transformed with lithium acetate method. All the mutants were confirmed by PCR, and DNA sequencing.

For the yeast spot sensitivity assays, yeast strains were grown overnight at 30 °C in YPD, diluted to an OD_600_ of 0.1, and grown for another 6 hours in YPD. Then 5-fold serial dilutions of cultures were spotted onto different media and grown at 30 °C for 3 days. To examine the utilization of various carbon sources, cells were grown on plates containing 2% glucose (YPD), 2% potassium acetate and 3% glycerol.

### HAT assays

In the HAT assay with competitor peptide, the reactions were performed in the MES buffer (25 mM MES, 180 mM KCl, 2 mM MgCl_2_, 0.1 mg/ml BSA, 50 µM acetyl-CoA, 0.5 µM NCP and 500 µM each competitor peptide, pH6.5). The peptide sequences used in this assay as follows: LANA (MAPPGMRLRSGRSTGAPLTRGS), R9E (MAPPGMRLESGRSTGAPLTRGS), Epl1_RA (ASAGSSNSRFRHRKISVKQHL), R56E (ASAGSSNSRFEHRKISVKQHL), R58E (ASAGSSNSRFRHEKISVKQHL).

The reactions were initiated by adding 10 nM NuA4. After incubation on ice for 25 min, the reactions were terminated by adding SDS loading buffer and heated at 100 °C for 1 min. The reaction mixtures were resolved with gradient SDS-PAGE, followed by Western blotting analysis using the primary antibody against pan-acetylated lysine (Cell Signaling Technology, Cat# 9681S). After exposure, the signals were quantified by ImageJ. Statistical analyses were performed using GraphPad Prism, and the statistical significance was determined using unpaired two-tailed Student’s *t*-test.

In the HAT assay with Gal4-VP16, NuA4 and Gal4-VP16 were premixed at a molar ratio of 1:1. The reactions were performed in the buffer (20 mM HEPES, 50 mM KCl, 2 mM MgCl_2_, 0.1 mg/ml BSA, 50 µM acetyl-CoA, 0.1 µM NCP and 1 µg salmon sperm DNA, pH 7.6). The reactions were initiated by adding 0.07 µM NuA4-Gal4-VP16 mixture or NuA4 alone, and proceeded at room temperature for 40 min. The following processes were the same as described above.

In the HAT assay with WT and DM NuA4, reactions were performed in a buffer containing 20 mM HEPES, 50 mM KCl, 2 mM MgCl_2_, 0.1 mg/ml BSA, 0.5 µM acetyl-CoA and 0.5 µM NCP, pH 7.6. The reactions were initiated by adding 10 nM WT or DM NuA4 and proceeded on ice for 10 min. The following processes were the same as described above.

## Acknowledgements

We thank Soo-Chen Cheng (Institute of Molecular Biology, Academia Sinica) for providing the yeast strain BJ2168, the Tsinghua University Branch of the China National Center for Protein Sciences (Beijing) for the cryo-EM facility. This work was supported by the National Key Research and Development Program (2019YFA0508902 to Z.C), the National Natural Science Foundation of China (32130016 and 31825016 to Z.C), Beijing Frontier Research Center for Biological Structure, Beijing Advanced Innovation Center for Structural Biology, and Tsinghua-Peking Joint Center for Life Sciences.

## Author Contribution

Q.K. prepared the sample, performed the biochemical analysis, and built the atomic model; K.C., W.H., and X.L. performed the EM analysis; Z.C. wrote the manuscript with help from all authors; Z.C. directed and supervised all of the research.

## Data Availability Statement

Coordinates and EM maps are deposited in the Protein Data Bank under accession codes EMD-32149, PDB ID 7VVY (TRA module); EMD-32148, PDB ID 7VVU (NCP-HAT); EMD-32150, PDB ID 7VVZ (NuA4-NCP); EMD-32156 (ARP submodule); EMD-32157 (Tra1 Head).

## References

1 Luger, K., Mader, A. W., Richmond, R. K., Sargent, D. F. & Richmond, T. J. Crystal structure of the nucleosome core particle at 2.8 A resolution. Nature 389, 251–260 (1997).

2 Kayne, P. S. et al. Extremely conserved histone H4 N terminus is dispensable for growth but essential for repressing the silent mating loci in yeast. Cell 55, 27–39 (1988).

3 Dorigo, B., Schalch, T., Bystricky, K. & Richmond, T. J. Chromatin fiber folding: requirement for the histone H4 N-terminal tail. J. Mol. Biol. 327, 85–96 (2003).

4 Shogren-Knaak, M. et al. Histone H4-K16 acetylation controls chromatin structure and protein interactions. Science 311, 844–847 (2006).

5 Allard, S. et al. NuA4, an essential transcription adaptor/histone H4 acetyltransferase complex containing Esa1p and the ATM-related cofactor Tra1p. EMBO J. 18, 5108–5119 (1999).

6 Bird, A. W. et al. Acetylation of histone H4 by Esa1 is required for DNA double-strand break repair. Nature 419, 411–415 (2002).

7 Babiarz, J. E., Halley, J. E. & Rine, J. Telomeric heterochromatin boundaries require NuA4-dependent acetylation of histone variant H2A.Z in Saccharomyces cerevisiae. Genes Dev. 20, 700–710 (2006).

8 Durant, M. & Pugh, B. F. NuA4-directed chromatin transactions throughout the Saccharomyces cerevisiae genome. Mol. Cell Biol. 27, 5327–5335 (2007).

9 Bruzzone, M. J., Grunberg, S., Kubik, S., Zentner, G. E. & Shore, D. Distinct patterns of histone acetyltransferase and Mediator deployment at yeast protein-coding genes. Genes Dev. 32, 1252–1265 (2018).

10 Dhar, S., Gursoy-Yuzugullu, O., Parasuram, R. & Price, B. D. The tale of a tail: histone H4 acetylation and the repair of DNA breaks. Philos. Trans. R. Soc. Lond. B Biol. Sci. 372(2017).

11 Doyon, Y. & Cote, J. The highly conserved and multifunctional NuA4 HAT complex. Curr. Opin. Genet. Dev. 14, 147–154 (2004).

12 Lee, K. K. & Workman, J. L. Histone acetyltransferase complexes: one size doesn’t fit all. Nat. Rev. Mol. Cell Biol. 8, 284–295 (2007).

13 Cheung, A. C. M. & Diaz-Santin, L. M. Share and share alike: the role of Tra1 from the SAGA and NuA4 coactivator complexes. Transcription 10, 37–43 (2019).

14 Boudreault, A. A. et al. Yeast enhancer of polycomb defines global Esa1-dependent acetylation of chromatin. Genes Dev. 17, 1415–1428 (2003).

15 Xu, P. et al. The NuA4 Core Complex Acetylates Nucleosomal Histone H4 through a Double Recognition Mechanism. Mol. Cell 63, 965–975 (2016).

16 Chittuluru, J. R. et al. Structure and nucleosome interaction of the yeast NuA4 and Piccolo-NuA4 histone acetyltransferase complexes. Nat. Struct. Mol. Biol. 18, 1196–1203 (2011).

17 Wang, X., Ahmad, S., Zhang, Z., Cote, J. & Cai, G. Architecture of the Saccharomyces cerevisiae NuA4/TIP60 complex. Nat. Commun. 9, 1147 (2018).

18 Setiaputra, D. et al. Molecular Architecture of the Essential Yeast Histone Acetyltransferase Complex NuA4 Redefines Its Multimodularity. Mol. Cell Biol. 38(2018).

19 Wang, H. et al. Structure of the transcription coactivator SAGA. Nature 577, 717–720 (2020).

20 Wu, J., Xie, N., Wu, Z., Zhang, Y. & Zheng, Y. G. Bisubstrate Inhibitors of the MYST HATs Esa1 and Tip60. Bioorg. Med. Chem. 17, 1381–1386 (2009).

21 Yuan, H. et al. MYST protein acetyltransferase activity requires active site lysine autoacetylation. EMBO J. 31, 58–70 (2012).

22 Sun, B. et al. Molecular basis of the interaction of Saccharomyces cerevisiae Eaf3 chromo domain with methylated H3K36. J. Biol. Chem. 283, 36504–36512 (2008).

23 Xu, C., Cui, G., Botuyan, M. V. & Mer, G. Structural basis for the recognition of methylated histone H3K36 by the Eaf3 subunit of histone deacetylase complex Rpd3S. Structure 16, 1740–1750 (2008).

24 Rossetto, D. et al. Eaf5/7/3 form a functionally independent NuA4 submodule linked to RNA polymerase II-coupled nucleosome recycling. EMBO J. 33, 1397–1415 (2014).

25 Lin, L., Chamberlain, L., Zhu, L. J. & Green, M. R. Analysis of Gal4-directed transcription activation using Tra1 mutants selectively defective for interaction with Gal4. Proc. Natl. Acad. Sci. USA 109, 1997–2002 (2012).

26 Knutson, B. A. & Hahn, S. Domains of Tra1 important for activator recruitment and transcription coactivator functions of SAGA and NuA4 complexes. Mol. Cell Biol. 31, 818–831 (2011).

27 Brown, C. E. et al. Recruitment of HAT complexes by direct activator interactions with the ATM-related Tra1 subunit. Science 292, 2333–2337 (2001).

28 Elias-Villalobos, A., Toullec, D., Faux, C., Seveno, M. & Helmlinger, D. Chaperone-mediated ordered assembly of the SAGA and NuA4 transcription co-activator complexes in yeast. Nat. Commun. 10, 5237 (2019).

29 Auger, A. et al. Eaf1 is the platform for NuA4 molecular assembly that evolutionarily links chromatin acetylation to ATP-dependent exchange of histone H2A variants. Mol. Cell Biol. 28, 2257–2270 (2008).

30 Bittner, C. B., Zeisig, D. T., Zeisig, B. B. & Slany, R. K. Direct physical and functional interaction of the NuA4 complex components Yaf9p and Swc4p. Eukaryot Cell 3, 976–983 (2004).

31 Klein, B. J. et al. Yaf9 subunit of the NuA4 and SWR1 complexes targets histone H3K27ac through its YEATS domain. Nucleic Acids Res. 46, 421–430 (2018).

32 Mitchell, L. et al. Functional dissection of the NuA4 histone acetyltransferase reveals its role as a genetic hub and that Eaf1 is essential for complex integrity. Mol. Cell Biol. 28, 2244–2256 (2008).

33 Downs, J. A. et al. Binding of chromatin-modifying activities to phosphorylated histone H2A at DNA damage sites. Mol. Cell 16, 979–990 (2004).

34 Turegun, B., Kast, D. J. & Dominguez, R. Subunit Rtt102 controls the conformation of the Arp7/9 heterodimer and its interactions with nucleotide and the catalytic subunit of SWI/SNF remodelers. J. Biol. Chem. 288, 35758–35768 (2013).

35 Schubert, H. L. et al. Structure of an actin-related subcomplex of the SWI/SNF chromatin remodeler. Proc. Natl. Acad. Sci. USA 110, 3345–3350 (2013).

36 Huang, J. & Tan, S. Piccolo NuA4-catalyzed acetylation of nucleosomal histones: critical roles of an Esa1 Tudor/chromo barrel loop and an Epl1 enhancer of polycomb A (EPcA) basic region. Mol. Cell Biol. 33, 159–169 (2013).

37 Barbera, A. J. et al. The nucleosomal surface as a docking station for Kaposi’s sarcoma herpesvirus LANA. Science 311, 856–861 (2006).

38 Steunou, A. L. et al. Combined Action of Histone Reader Modules Regulates NuA4 Local Acetyltransferase Function but Not Its Recruitment on the Genome. Mol. Cell Biol. 36, 2768–2781 (2016).

## References

39 Ashkenazy, H. et al. ConSurf 2016: an improved methodology to estimate and visualize evolutionary conservation in macromolecules. Nucleic Acids Res. 44, W344–350 (2016).

40 Gietz, R. D. & Schiestl, R. H. High-efficiency yeast transformation using the LiAc/SS carrier DNA/PEG method. Nat Protoc 2, 31–34 (2007).

41 Stark, H. GraFix: stabilization of fragile macromolecular complexes for single particle cryo-EM. Methods Enzymol. 481, 109–126 (2010).

42 Simon, M. D. et al. The site-specific installation of methyl-lysine analogs into recombinant histones. Cell 128, 1003–1012 (2007).

43 Dawson, P. E., Muir, T. W., Clark-Lewis, I. & Kent, S. B. Synthesis of proteins by native chemical ligation. Science 266, 776–779 (1994).

44 Li, Y. T. et al. A semisynthetic Atg3 reveals that acetylation promotes Atg3 membrane binding and Atg8 lipidation. Nat. Commun. 8, 14846 (2017).

45 Dyer, P. N. et al. Reconstitution of nucleosome core particles from recombinant histones and DNA. Methods Enzymol. 375, 23–44 (2004).

46 Mindell, J. A. & Grigorieff, N. Accurate determination of local defocus and specimen tilt in electron microscopy. J. Struct. Biol. 142, 334–347 (2003).

47 Ludtke, S. J. 3-D structures of macromolecules using single-particle analysis in EMAN. Methods Mol. Biol. 673, 157–173 (2010).

48 Scheres, S. H. RELION: implementation of a Bayesian approach to cryo-EM structure determination. J. Struct. Biol. 180, 519–530 (2012).

49 Frank, J. et al. SPIDER and WEB: processing and visualization of images in 3D electron microscopy and related fields. J. Struct. Biol. 116, 190–199 (1996).

50 Lei, J. & Frank, J. Automated acquisition of cryo-electron micrographs for single particle reconstruction on an FEI Tecnai electron microscope. J. Struct. Biol. 150, 69–80 (2005).

51 Zheng, S. Q. et al. MotionCor2: anisotropic correction of beam-induced motion for improved cryo-electron microscopy. Nat. Methods 14, 331–332 (2017).

52 Rohou, A. & Grigorieff, N. CTFFIND4: Fast and accurate defocus estimation from electron micrographs. J. Struct. Biol. 192, 216–221 (2015).

53 Nakane, T., Kimanius, D., Lindahl, E. & Scheres, S. H. Characterisation of molecular motions in cryo-EM single-particle data by multi-body refinement in RELION. Elife 7(2018).

54 Li, X., Zheng, S., Agard, D. A. & Cheng, Y. Asynchronous data acquisition and on-the-fly analysis of dose fractionated cryoEM images by UCSFImage. J. Struct. Biol. 192, 174–178 (2015).

55 Diaz-Santin, L. M., Lukoyanova, N., Aciyan, E. & Cheung, A. C. Cryo-EM structure of the SAGA and NuA4 coactivator subunit Tra1 at 3.7 angstrom resolution. Elife 6(2017).

56 Afonine, P. V. et al. Towards automated crystallographic structure refinement with phenix.refine. Acta Crystallogr. D Biol. Crystallogr. 68, 352–367 (2012).

57 Ye, Y. et al. Structure of the RSC complex bound to the nucleosome. Science 366, 838–843 (2019).

58 Graham, M., Combe, C. W., Kolbowski, L. & Rappsilber, J. xiView: A common platform for the downstream analysis of Crosslinking Mass Spectrometry data. bioRxiv (2019).

59 Pettersen, E. F. et al. UCSF ChimeraX: Structure visualization for researchers, educators, and developers. Protein Sci. 30, 70–82 (2021).

