## Extended Data Figures for "Structure of the NuA4 acetyltransferase complex bound to the nucleosome"

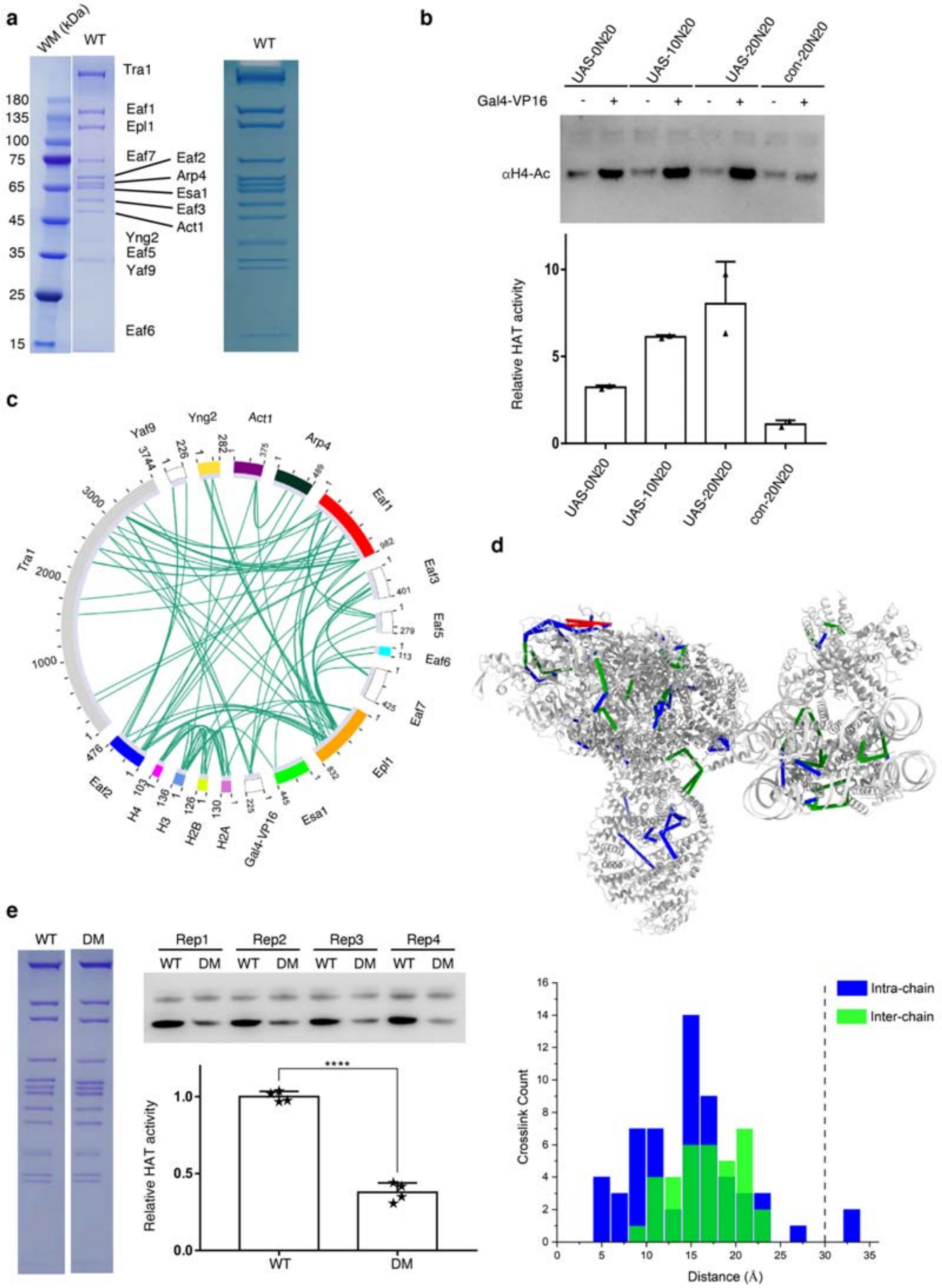


**Extended Data Fig. 1 Biochemistry of NuA4 and the crosslinking data detected by CL-MS.** (**a**) Preparation of endogenous NuA4 complex from *S.cerevisiae*. SDS-PAGE gels were showed with Coomassie stain. More sample of the WT complex was loaded to show the weak bands on the right panel. (**b**) Relative HAT activity of NuA4 towards the nucleosomes in the absence and presence of Gal4-VP16. The Gal4 binding site (UAS) was placed at different positions of the nucleosomes (UAS-0N20, UAS-10N20 and UAS-20N20), and the unrelated sequence was used as a control (con-20N20). Acetylation of H4 was detected by Western blotting. A presentative gel was shown on the top, and quantification at the bottom. The activities were normalized to the ones in the absence of Gal4-VP16. Error bars indicate SD (n = 2 technical replicates). (**c**) Overview of the cross-linking data. Circular plot of high confidence lysine-lysine inter-subunit crosslinks obtained by mass spectrometry for the NuA4-NCP complex. (**d**) Validated cross-links mapped onto the NuA4-NCP structure. Blue lines, the intra-chain cross-links with cross-linked sites within the 30 Å distance permitted by BS3; green lines, the inter-chain cross-links, whereas red lines depict cross-link over more than 30 Å. Bottom graph displays the distribution of cross-link distances. A vertical line indicates a cutoff at 30 Å, the distance considered reasonable for BS3 crosslinks. (**e**) Relative HAT activity of an independent batch of WT and DM mutant NuA4 different from that of Fig. 2f. Error bars indicate SD (n = 4 technical replicates). ****, $p<0.0001$.

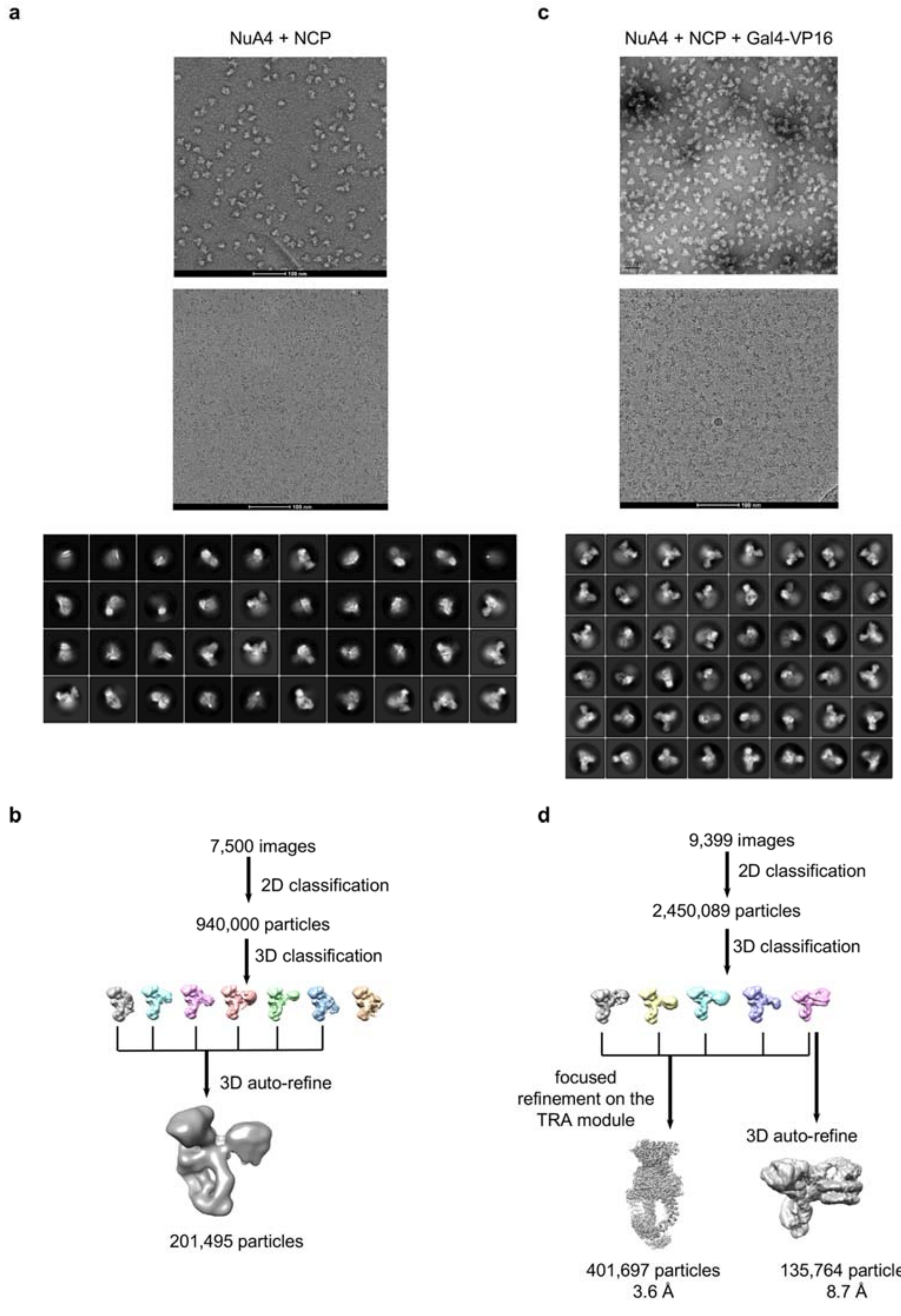


**Extended Data Fig. 2 Cryo-EM analysis of the NuA4-NCP complex in the absence and presence of Gal4-VP16**

(**a**) Representative negative stain images (top panel), cryo-EM images (middle panel), and 2D classification (bottom panel) of NuA4+NCP. (**b**) Flowcharts of the cryo-EM data processing for the dataset of NuA4+NCP. (**c**) Representative negative stain images (top panel), cryo-EM images (middle panel), and 2D classification (bottom panel) of NuA4+NCP+Gal4-VP16. (**d**) Flowcharts of the cryo-EM data processing for the dataset of NuA4+NCP+Gal4-VP16.

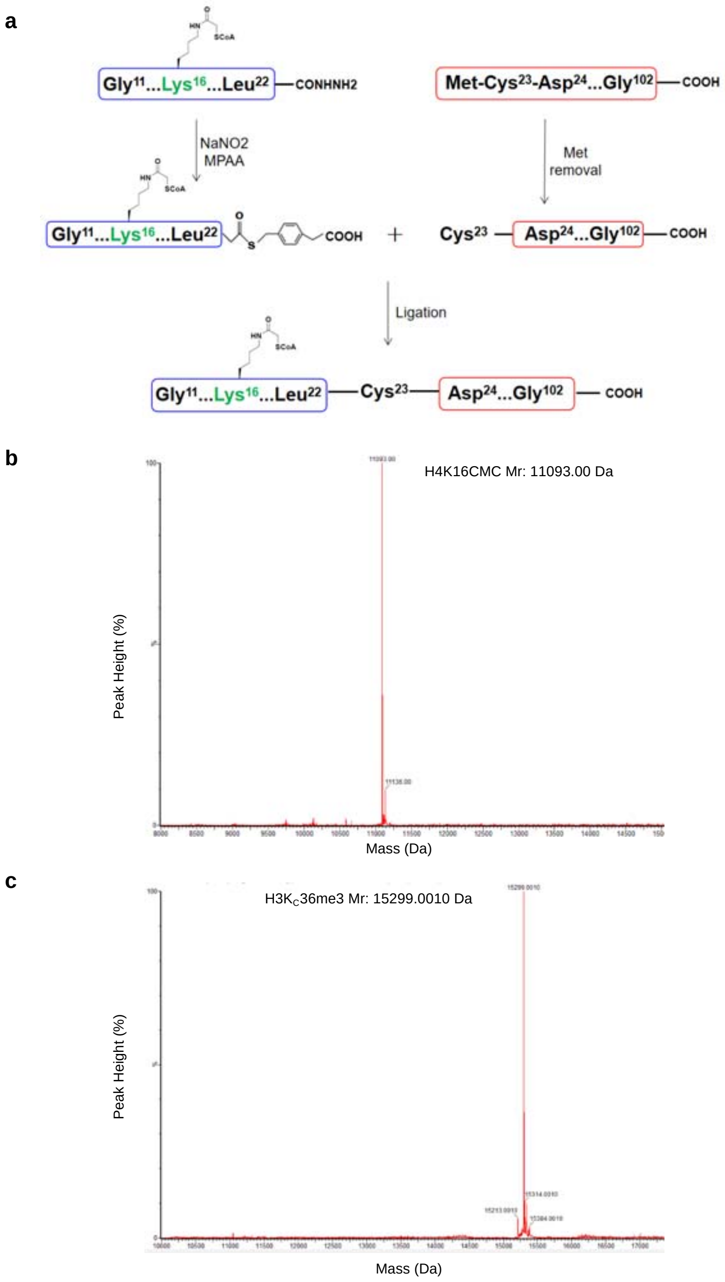


**Extended Data Fig. 3 Chemistry and Mass-spectrometry analyses of the modified histones**

(**a**) Schematic diagram of the chemical semi-synthesis of histone H4 with CMC modification at Lys16. (**b**) Mass-spectrometry identification of the CMC-modified H4 (the expected mass is 11092.57 Da). (**c**) Mass-spectrometry identification of the H3K$_C$36 me3-modified H3 (the expected mass is 15299.76 Da).

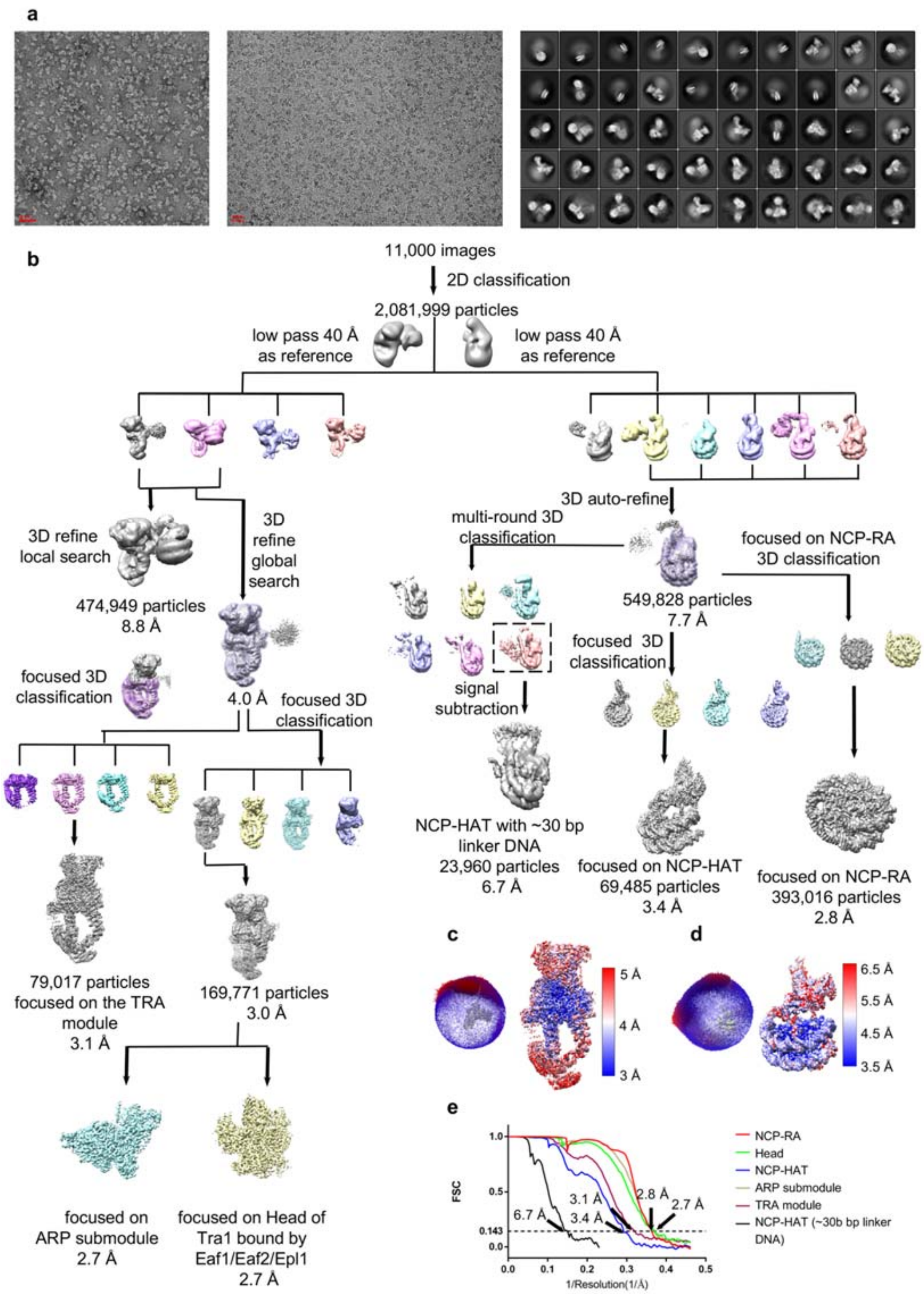


**Extended Data Fig. 4 Cryo-EM analysis of the NuA4-NCP complex in the presence of Gal4-VP16 and histone-modified nucleosome.**

(**a**) Representative negative stain images (left panel), cryo-EM images (middle panel), and 2D classification (right panel) of NuA4+Gal4-VP16 bound to the H4- and H3-modified nucleosome. (**b**) Flowcharts of the cryo-EM data processing for the dataset of NuA4+Gal4-VP16 bound to the H4- and H3-modified nucleosome. (**c-d**) Angular distributions of cryo-EM particles in the final round of refinement of the masked dataset and the estimation of the local resolutions of the TRA module (**c**) and the nucleosome bound HAT module (**d**). (**e**) Gold standard Fourier shell correlation (FSC) curves, showing the overall nominal resolutions of 6.7 Å, 3.4 Å, 3.1 Å, 2.8 Å, 2.7 Å and 2.7 Å for the NCP-HAT with longer linker DNA, the nucleosome bound HAT module, TRA module, the NCP bound with arginine anchors (RAs), ARP submodule, head of Tra1 bound with Eaf1/Eaf2/Epl1, respectively.

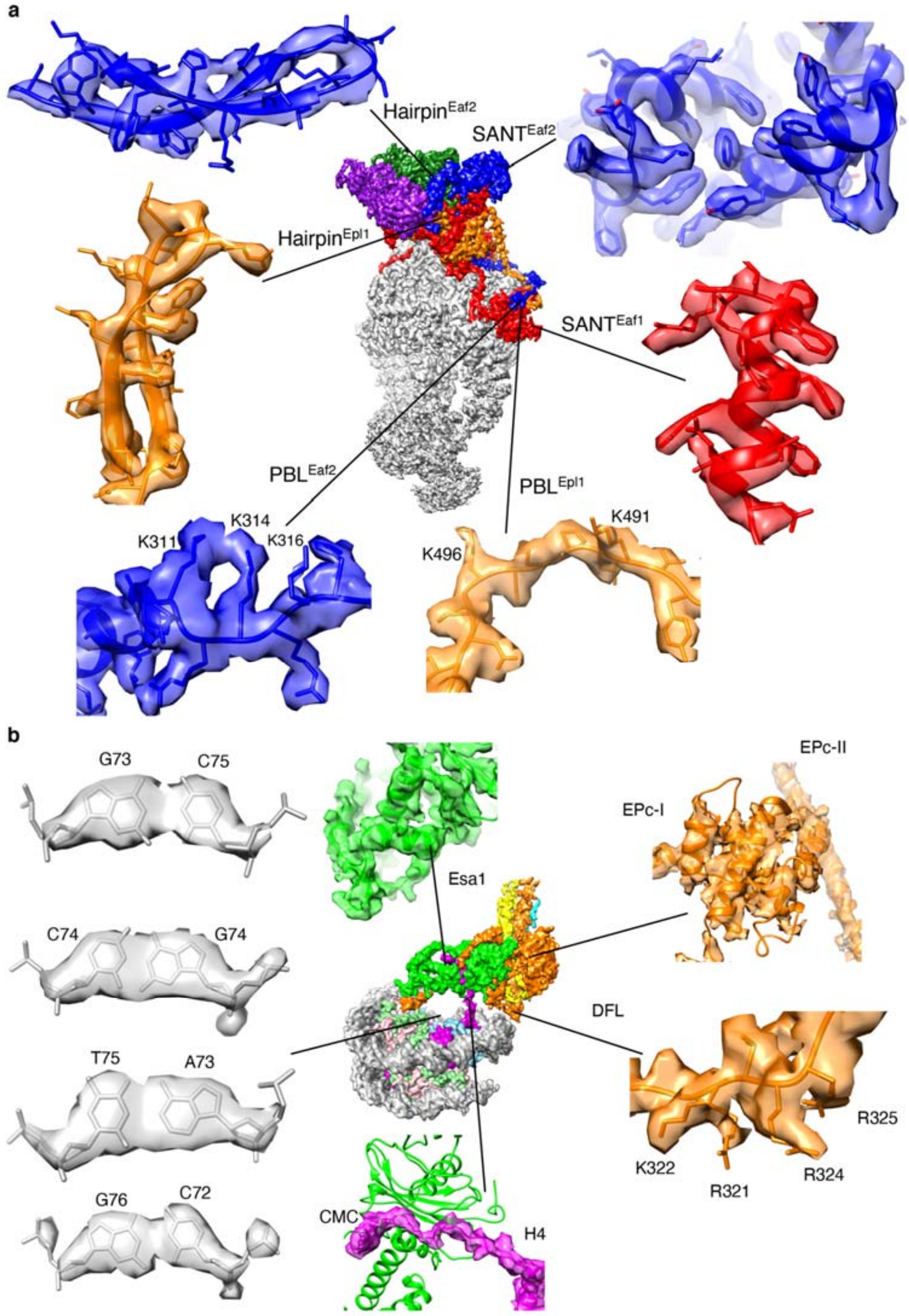


**Extended Data Fig. 5 Local density maps of the NuA4-nucleosome complex.** (**a**) Local cryo-EM maps of the TRA module. PBL, polybasic loop. (**b**) Local cryo-EM maps of the HAT module bound to the nucleosome. EM densities of four consecutive base pairs at the central position of the nucleosome are showed on the left.

```
ScEaf2      -MSSSDIFDVLNIKQKSRSPTNGQVS----------VPSSSAANRPKPQVTGMQRELFNL  49
HsDMAP1     MATGADVRDILELGGPEGDAASGTISKKDIINPDKKKSKKSSETLTFKRPEGMHREVYAL  60
MmDMAP1     MATGADVRDILELGGPEGDAASGTISKKDIINPDKKKSKKSSETLTFKRPEGMHREVYAL  60
DmDMAP1     --MSADVRDILDMERANTPE----VTRDSFLATKK--RNFERTKTASRRPEGMHREVFAL  52
                 *  *                                          ** **   *

ScEaf2      LGENQ----PPVVI-------KSGNNF-KEKMLSTSKPSPWSFVEFKAN------NSVTL  91
HsDMAP1     LYSDK-KDAPPLLPSDT----GQGYRTVKAKLG-SKKVRPWKWMPFTNPARK---DGAMF  111
MmDMAP1     LYSDK-KDAPPLLPSDT----GQGYRTVKAKLG-SKKVRPWKWMPFTNPARK---DGAMF  111
DmDMAP1     LYTDK-KDAPPLLPTDTALGIGAGYKETKARLG-MKKVRKWEWAPFSNPARN---DSAVF  107
            *        **                  *       *   *    *

ScEaf2      RHWVKGSKELIGDT---PKESPYSKFNQHLSIPSFTKE-EYEAFMNENEGTQKSVES---  144
HsDMAP1     FHWRRAAEE--------GKDYPFARFNKTVQVPVYSEQ-EYQLYLH--------------  148
MmDMAP1     FHWRRAAEE--------GKDYPFARFNKTVQVPVYSEQ-EYQLYLH--------------  148
DmDMAP1     HHWKRVTDN--------STDYPFAKFNKQLEVPSYTMT-EYNAHLRNN------------  146
             **                       **     *      **

ScEaf2      --------EKNHN-------ENFTNEKKDESKNSWSFEEIEYLFNLCKKYDLRWFLIFDR  189
HsDMAP1     -------------------------------DDAWTKAETDHLFDLSRRFDLRFVVIHDR  177
MmDMAP1     -------------------------------DDAWTKAETDHLFDLSRRFDLRFVVIHDR  177
DmDMAP1     -------------------------------INNWSKVQTDHLFDLARRFDLRFIVMADR  175
                                              *       ** *    ***     **

ScEaf2      YSYNN-S--RTLEDLKEKFYYTCRNYFKASDPSN-------PLL-SSLNFSAEKEIERKK  238
HsDMAP1     YDHQQF-KKRSVEDLKERYYHICAKLANVRAVPG--------TDLKIPVFDAGHERRRKE  228
MmDMAP1     YDHQQF-KKRSVEDLKERYYHICAKLANVRAVPG--------TDLKIPVFDAGHERRRKE  228
DmDMAP1     WNRQQH-GTKTVEELKERYYEVVALLAKAKNQ-T--------SEKKVFVYDVEHERRRKE  225
                        * ***  *                                  *  **

ScEaf2      YLQRLLSRSAAEIAEEEALVVESKKFEMAAKRTLAERESLLRLLDSPHSDQTITQYLTSQ  298
HsDMAP1     QLERLYNRTPEQVAEEEYLLQELRKIEARKKEREKRSQDLQKLITAADTTAE--QRRT--  284
MmDMAP1     QLERLYNRTPEQVAEEEYLLQELRKIEARKKEREKRSQDLQKLITAADTTAE--QRRT--  284
DmDMAP1     QLEKLFKRTTQQVEEENMLINEMKKIEARKKERERKTQDLQKLISQADQQNE--HASNT-  282
             *  *  *      **  *  *  * *   *        *  *

                               PBL
ScEaf2      GMSQLYNALLA-DKTRKRKHD------------LNIPENPW-----MKQQQQFAQHRQLQ  340
HsDMAP1     ------E-----RKAPKKKLP---------------------------------------  294
MmDMAP1     ------E-----RKAPKKKLP---------------------------------------  294
DmDMAP1     PSTRKYE-----KKLHKKKVH---------------------------------------  298
                         *  * *
```

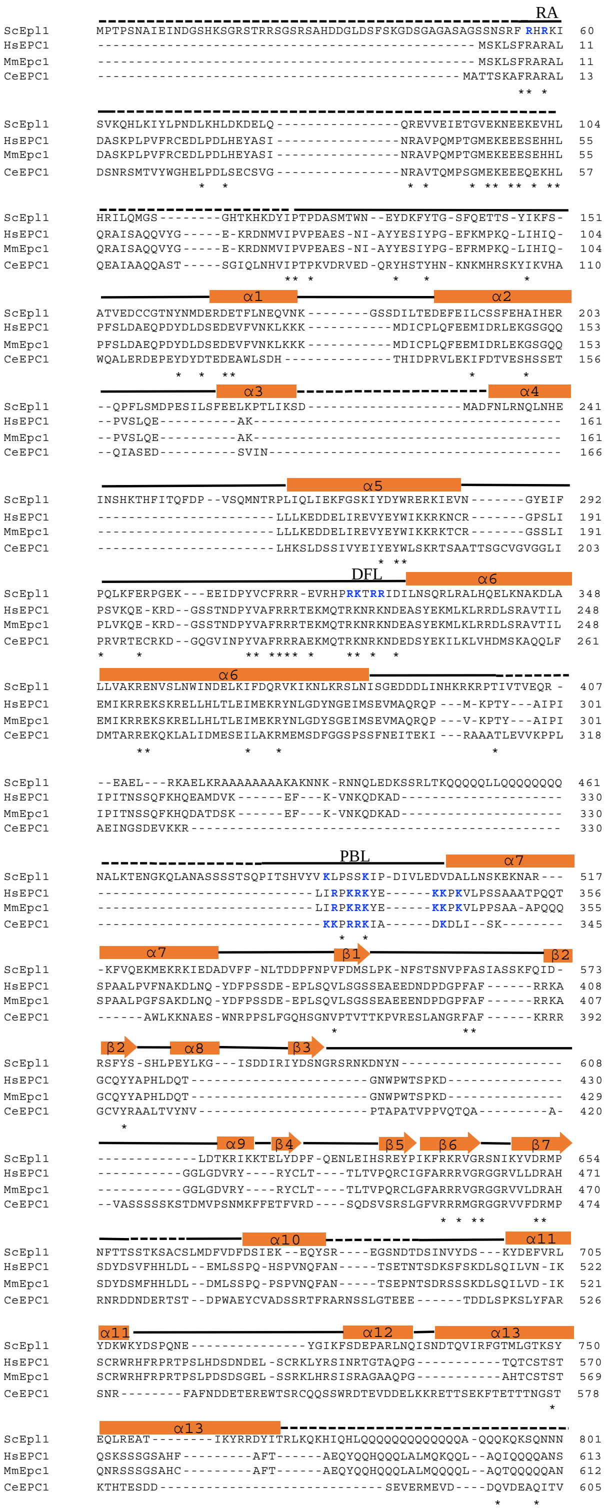


**Extended Data Fig. 7 Multiple sequence alignments of Epl1-like proteins.** The conserved residues are indicated by “*”. The secondary structural assignments are based on the structure of Epl1.

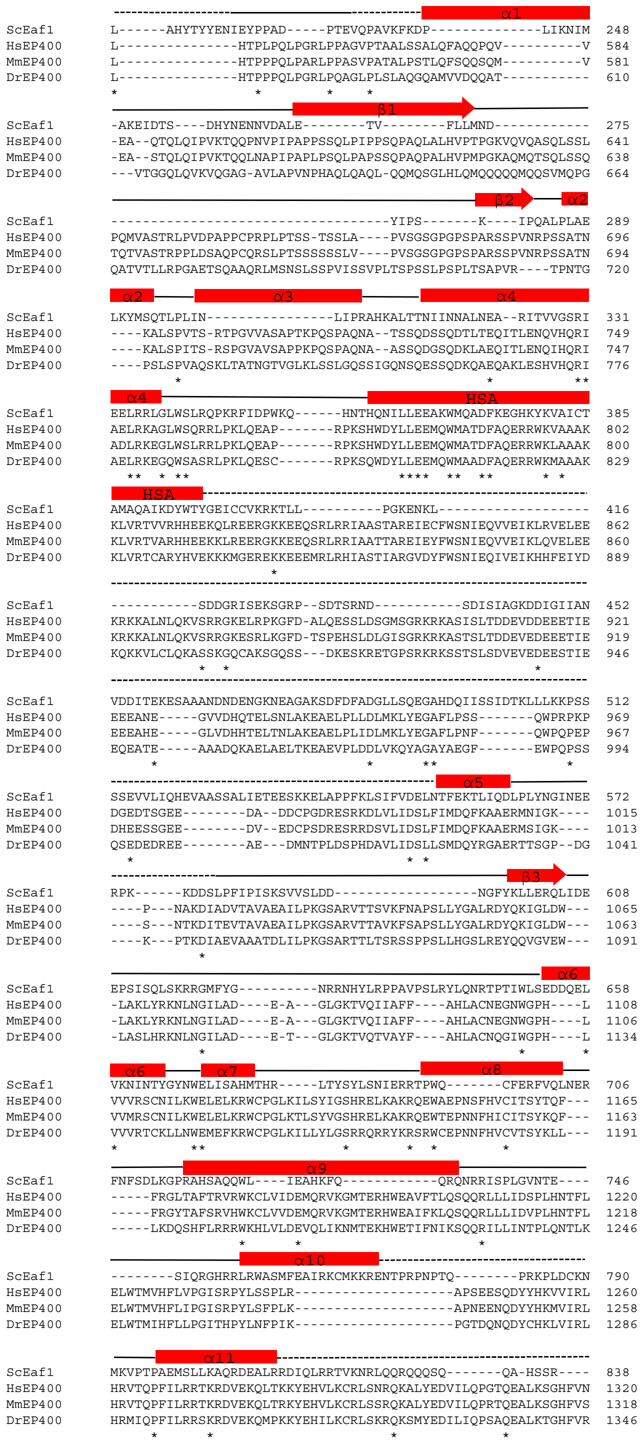


**Extended Data Fig. 8 Multiple sequence alignments of Eaf1-like proteins.** The conserved residues are indicated by "*". The secondary structural assignments are based on the structure of Eaf1.

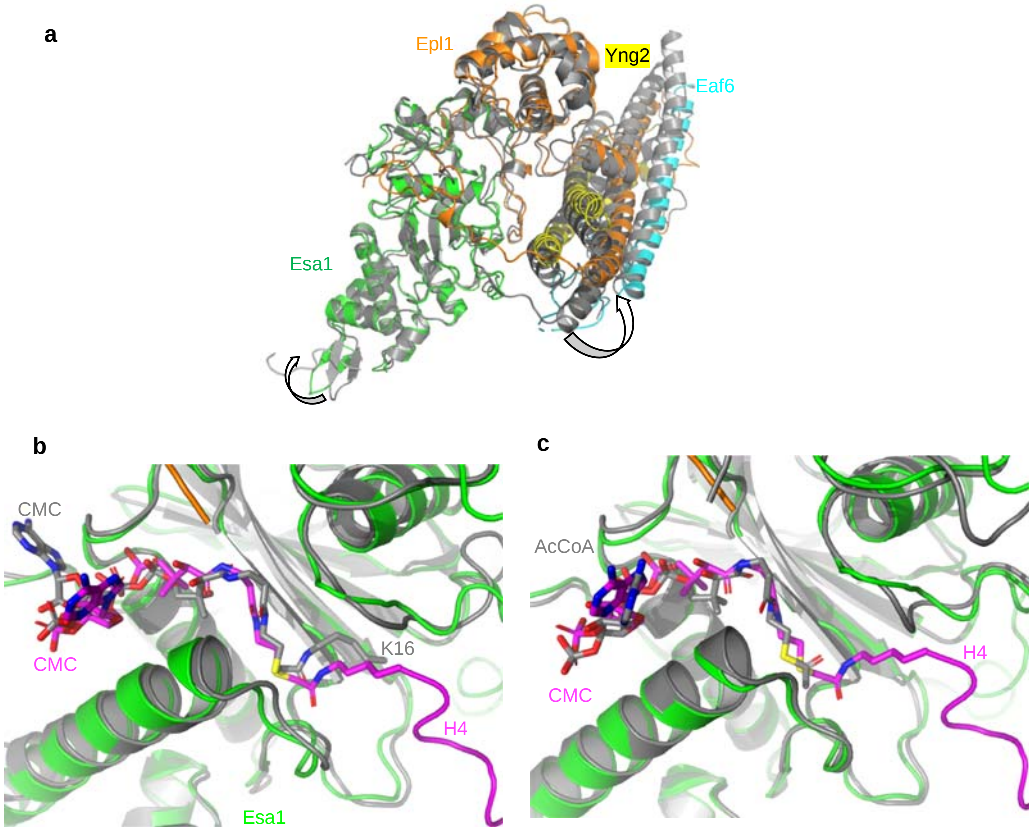


**Extended Data Fig. 9 Additional structural analysis of the HAT module.** (**a**) Comparison of the structures of the HAT module bound to the nucleosome (color coded) and the nucleosome-free Piccolo subcomplex (colored grey, PBD code 5J9W)[1]. The structure of the Esa1 subunit is aligned. (**b**) Structural alignment of CMC-binding pocket in NuA4 (color coded) and in Esa1 (colored grey, PBD code 3TO6)[2]. (**c**) Structural comparison of CMC-binding in NuA4 (color coded) and AcCoA in Piccolo (colored grey, PBD code 5J9W)[1].

1 Xu, P. *et al.* The NuA4 Core Complex Acetylates Nucleosomal Histone H4 through a Double Recognition Mechanism. Mol. Cell 63, 965-975 (2016).
2 Yuan, H. *et al.* MYST protein acetyltransferase activity requires active site lysine autoacetylation. EMBO J. 31, 58-70 (2012).
