## Extended Data Table 1 for "Structure of the NuA4 acetyltransferase complex bound to the nucleosome"

**Extended Data Table 1 Cryo-EM data collection, refinement and validation statistics**

| | #1 TRA module (EMD-32149, PDB 7VVY) | #2 NCP-HAT (EMD-32148, PDB 7VVU) | #3 NuA4-NCP (EMD-32150, PDB 7VVZ) | #4 ARP submodule (EMD-32156) | #5 Tra1 Head (EMD-32157) | #6 NCP-RA (EMD-32158) | #7 NCP-HAT with long linker (EMD-33224) |
|---|---|---|---|---|---|---|---|
| **Data collection and processing** | | | | | | | |
| Microscope | Krios G3i | Krios G3i | Krios G3i | Krios G3i | Krios G3i | Krios G3i | Krios G3i |
| Camera | K3 | K3 | K3 | K3 | K3 | K3 | K3 |
| Magnification (nominal) | 81,000 | 81,000 | 81,000 | 81,000 | 81,000 | 81,000 | 81,000 |
| Electron exposure ($e^-/Å^2$) | 50 | 50 | 50 | 50 | 50 | 50 | 50 |
| Number of frames collected | 32 | 32 | 32 | 32 | 32 | 32 | 32 |
| Energy filter slit width (eV) | 20 | 20 | 20 | 20 | 20 | 20 | 20 |
| Automation software | AutoEMation2 | AutoEMation2 | AutoEMation2 | AutoEMation2 | AutoEMation2 | AutoEMation2 | AutoEMation2 |
| Voltage (kV) | 300 | 300 | 300 | 300 | 300 | 300 | 300 |
| Micrographs (no.) | 11,000 | 11,000 | 11,000 | 11,000 | 11,000 | 11,000 | 11,000 |
| Defocus range (μm) | -1.3— -1.8 | -1.3— -1.8 | -1.3— -1.8 | -1.3— -1.8 | -1.3— -1.8 | -1.3— -1.8 | -1.3— -1.8 |
| Pixel size (Å) | 0.54125 | 0.54125 | 0.54125 | 0.54125 | 0.54125 | 0.54125 | 0.54125 |
| Symmetry imposed | C1 | C1 | C1 | C1 | C1 | C1 | C1 |
| Initial particle images (no.) | 2,081,999 | 2,081,999 | 2,081,999 | 2,081,999 | 2,081,999 | 2,081,999 | 2,081,999 |
| Final particle images (no.) | 79,017 | 69,485 | 474,949 | 169,771 | 169,771 | 393,016 | 23,960 |
| Error of translations | 0.462 | 0.958 | 0.469 | 0.75 | 0.859 | 0.601 | 0.949 |
| Error of rotations | 0.697 | 0.744 | 2.607 | 0.493 | 0.407 | 0.83 | 2.299 |
| Map resolution (Å) (masked) | 3.1 | 3.4 | 8.8 | 2.7 | 2.8 | 2.8 | 6.7 |
| FSC threshold | 0.143 | 0.143 | 0.143 | 0.143 | 0.143 | 0.143 | 0.143 |
| Map sharpening *B* factor ($Å^2$) | -20 | -20 | | -60 | -40 | -30 | -30 |
| **Refinement** | | | | | | | |
| Initial model used (PDB code) | 5I9E, 5OJS | 5Z3V, 5J9W, 3TO6 | | | | | |
| Refinement package | Phenix | Phenix | | | | | |
| Model-map scores | | | | | | | |
| CC(mask) | 0.79 | 0.67 | | | | | |
| CC(box) | 0.80 | 0.79 | | | | | |
| CC(peaks) | 0.71 | 0.66 | | | | | |
| CC(volume) | 0.78 | 0.72 | | | | | |

| | | |
|---|---|---|
| R.m.s. deviations | | |
| Bond lengths (Å) | 0.013 | 0.005 |
| Bond angles (°) | 0.971 | 0.654 |
| C-beta deviation | 0.00 | 0.00 |
| EMRinger score | 1.78 | 1.79 |
| CaBLAM outliers | 4.89 | 1.20 |
| **Validation** | | |
| MolProbity score | 1.91 | 1.47 |
| Clashscore | 8.15 | 8.78 |
| Poor rotamers (%) | 0.48 | 0.31 |
| Ramachandran plot | | |
| Favored (%) | 92.00 | 98.47 |
| Allowed (%) | 7.90 | 1.53 |
| Disallowed (%) | 0.10 | 0.00 |
